# Increased LPS-Induced Fever and Sickness Behavior in Adult Male and Female Rats Perinatally Exposed to Morphine

**DOI:** 10.1101/2023.09.20.558690

**Authors:** Hannah J. Harder, Morgan G. Gomez, Christopher T. Searles, Meghan E. Vogt, Anne Z. Murphy

**Author notes:** Corresponding Author: Anne Z. Murphy, 100 Piedmont Ave., Atlanta, GA, 30303.

## Abstract

As a result of the current opioid crisis, the rate of children born exposed to opioids has skyrocketed. Later in life, these children have an increased risk for hospitalization and infection, raising concerns about potential immunocompromise, as is common with chronic opioid use. Opioids can act directly on immune cells or indirectly via the central nervous system to decrease immune system activity, leading to increased susceptibility, morbidity, and mortality to infection. However, it is currently unknown how perinatal opioid exposure (POE) alters immune function. Using a clinically relevant and translatable model of POE, we have investigated how baseline immune function and the reaction to an immune stimulator, lipopolysaccharide, is influenced by *in utero* opioid exposure in adult male and female rats. We report here that POE potentiates the febrile and neuroinflammatory response to lipopolysaccharide, likely as a consequence of suppressed immune function at baseline (including reduced antibody production). This suggests that POE increases susceptibility to infection by manipulating immune system development, consistent with the clinical literature. Investigation of the mechanisms whereby POE increases susceptibility to pathogens is critical for the development of potential interventions for immunosuppressed children exposed to opioids *in utero*.

**Highlights:**

- Perinatal exposure to morphine leads to increased fever and sickness post-LPS
- Perinatal exposure to morphine alters peripheral immunity
- Perinatal exposure to morphine increases microglial activation

## 1. Introduction

The exponential increase in opioid use in the United States, particularly among women of reproductive age, has resulted in a surge of infants exposed to opioids *in utero*. Most of these infants will experience opioid withdrawal at birth (neonatal opioid withdrawal syndrome; NOWS), requiring an extended stay in the neonatal intensive care unit (Kocherlakota, 2014). Gestation and the early postnatal period are critical periods of immune system development in humans and rodents (Georgountzou and Papadopoulos, 2017); however, little is known about the long-term consequences of perinatal opioid exposure on immune function. In adults, chronic opioid use is associated with suppression of the peripheral immune system, including decreased natural killer cell cytotoxicity (Beilin et al., 1996, 1992, 1989; Fecho and Lysle, 1999; Nelson et al., 2000; Novick et al., 1989; Sacerdote et al., 1997; Yokota et al., 2004), reduced macrophage phagocytosis (Casellas et al., 1991; Lugo-Chinchilla et al., 2006; Tomassini et al., 2004; Tomei and Renaud, 1997), and altered proinflammatory cytokine production (Clark et al., 2007; Madera-Salcedo et al., 2011; Stoll-Keller et al., 1997; Wang et al., 2011). Clinical chart review suggests that infants exposed to opioids *in utero* are similarly at an increased risk of infection and rehospitalization (Arter et al., 2021; Uebel et al., 2015; Witt et al., 2017), suggesting parallel suppression of immune function. Chronic opioid-induced deficits in antibody production are of particular concern to opioid-exposed infants (Bussiere et al., 1992; Eisenstein et al., 1993; Taub et al., 1991), as this response is an essential component of the adaptive immune system and serves to form immunological memories of previous exposure. Indeed, deficits in antibody production increase susceptibility to infection, as the adaptive immune system is unable to recognize and eliminate pathogens (Barmettler et al., 2018).

The periaqueductal gray (PAG), a critical neural substrate in opioid signaling (Loyd et al., 2008), has been implicated in the central immunosuppressive effects of opioids (Gomez-Flores and Weber, 2000). Direct administration of morphine into the PAG suppresses a number of immune cell functions, including natural killer cell cytotoxic activity, lymphocyte proliferation, and cytokine production (Gomez-Flores et al., 1999; Weber and Pert, 1989). Furthermore, morphine has a stimulatory effect on PAG microglia to induce cytokine release in a Toll-like receptor 4 (TLR4)-dependent manner (Bokhari et al., 2009; Eidson and Murphy, 2013; Lee et al., 2018; Zhang et al., 2020, 2011). Morphine has also been shown to suppress peripheral immune cell activity in a TLR4-dependent manner (Zhang et al., 2020). Therefore, morphine is poised to impact the immune system at multiple levels, promoting neuroinflammation centrally, and immunosuppression and increased susceptibility to pathogens peripherally.

Systemic administration of lipopolysaccharide (LPS), which mimics a Gram-negative bacterial infection, is one of the most common experimental immune models (Lasselin et al., 2020). LPS is a pathogen-associated molecular pattern (PAMP) that binds primarily to the innate immune receptor TLR4, leading to peripheral cytokine production (Zampronio et al., 2015). These cytokines act primarily within the hypothalamic median preoptic area to induce prostaglandin E2 synthesis, promoting fever via increased brown adipose tissue metabolism and vasoconstriction (Hart, 1988; Machado et al., 2020; Saper and Breder, 1994; Zampronio et al., 2015). LPS-induced fever is accompanied by a characteristic set of behaviors associated with sickness, including anorexia, lethargy, and reduced grooming (Hart, 1988). Both fever and sickness serve to engage the immune system to restrict pathogen growth and reduce metabolic demand to facilitate fever (Hart, 1988); therefore, any alterations in the course of a typical fever and sickness response may increase the severity of infection. Counterintuitively, immunosuppression generally leads to elevated febrile response to LPS (Miñano et al., 2004; Tavares et al., 2006, 2005) or infection (Oude Nijhuis et al., 2002) and is often the only sign of infection in immunosuppressed patients (Pizzo, 1999). Although the mechanism by which immunosuppression augments infection-induced fevers is currently unknown, it is thought that immunosuppression leads to an inability of the body to properly mount a cytokine response to infection, leading to an elevated, centrally-mediated fever response (Oude Nijhuis et al., 2002).

Limited preclinical evidence has associated perinatal opioid exposure with immune dysregulation after LPS exposure (Hamilton et al., 2007; Shavit et al., 1998). Previous studies examining the impact of perinatal opioid exposure on immune function utilized dosing paradigms that fail to mirror the clinical profile of women who use opioids while pregnant. The present studies were conducted to address this gap using a clinically relevant and translatable model of perinatal opioid exposure (POE). We hypothesize that perinatal exposure to opioids results in immune system dysregulation, both centrally and peripherally, leading to an increased immune response to LPS.

## 2. Methods

### 2.1 Experimental subjects

All experiments utilized male and female Sprague Dawley rats (Charles River Laboratories, Boston, MA). Rats were housed in same-sex pairs or groups of three on a 12:12 hours light/dark cycle (lights on at 8:00 AM) in Optirat GenII individually ventilated cages (Animal Care Systems, Centennial, Colorado, USA) with corncob bedding. Food (Lab Diet 5001 or Lab Diet 5015 for breeding pairs, St. Louis, MO, USA) and water were provided *ad libitum* throughout the experiment, except during testing. These studies were approved by the Institutional Animal Care and Use Committee at Georgia State University and performed in compliance with the National Institutes of Health Guide for the Care and Use of Laboratory Animals. Every effort was made to reduce the number of rats used and minimize pain and suffering.

### 2.2 Perinatal opioid exposure paradigm

Female Sprague Dawley rats (P60) were implanted with iPrecio® SMP-200 microinfusion minipumps under isoflurane anesthesia (Harder et al., 2023). See Figure 1 for a description of the dosing paradigm. Pumps were programmed to deliver morphine at 10 mg/kg three times a day. One week after morphine initiation, females were paired with sexually-experienced males for two weeks to induce pregnancy. Morphine exposure to the dams continued throughout gestation, with doses increasing weekly by 2 mg/kg until 16 mg/kg was reached. Dams continued to receive morphine after parturition, such that pups received morphine indirectly. Beginning at P5, morphine dosage was decreased by 2 mg/kg daily until P7, when the dose reached 0 mg/kg. Control rats were implanted with pumps filled with sterile saline. No differences were noted in maternal behavior of morphine vs. vehicle dams (Harder et al., 2023). Pups were weaned at P21 into treatment-matched cages, where they remained until adulthood.

**Figure 1.**
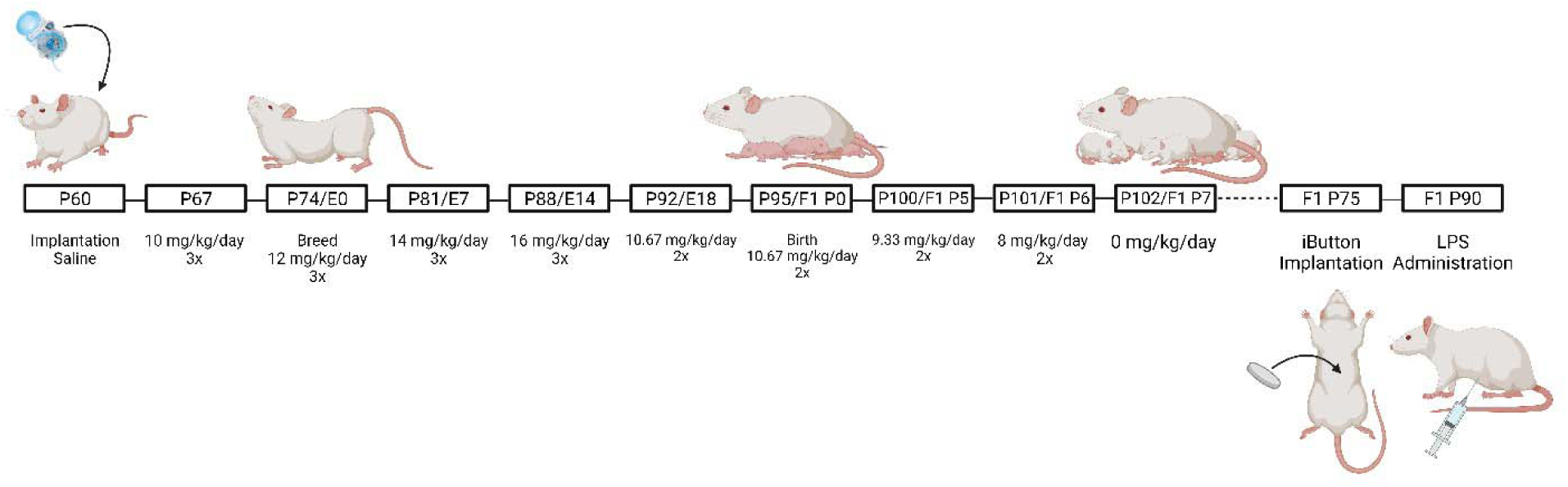
Schematic of the perinatal opioid exposure dosing paradigm. Created with Biorender.

### 2.3 iButton implantation

At P75, male and female rats born to mothers exposed to morphine (MOR) or vehicle (VEH) were anesthetized with 5% isoflurane and maintained at 2- 3%. A midline abdominal incision was made using sterile surgical techniques, and a wax-coated iButton temperature logger (Thermochron DS1922L) was placed into the abdominal cavity. All rats received carprofen (5 mg/mL/kg; i.p.) prior to and twenty-four hours post-surgery for pain relief. iButton loggers were programmed to record core body temperature in 10-minute intervals beginning twenty-four hours before and twenty-four post-LPS administration.

### 2.4 Lipopolysaccharide treatment

Approximately 14 days following iButton implantation, rats were administered lipopolysaccharide (250 ug/kg/mL, i.p.) derived from *E. coli* O111:B4 (Sigma-Aldrich; L2630) to induce fever and sickness. Body temperature was recorded for 24 hours post-LPS. Every two hours, LPS-induced sickness behavior was quantified using an 8-point Likert scale, which measured changes in physical features (ear changes, nose/cheek flattening, orbital tightening, and piloerection), modified from (Sotocinal et al., 2011). See Figure 2 for an example of sickness-associated features. Rats received a score of 0 if absent, 1 if present, and 2 if present and severe, for a maximum sickness score of 8.

**Figure 2.**
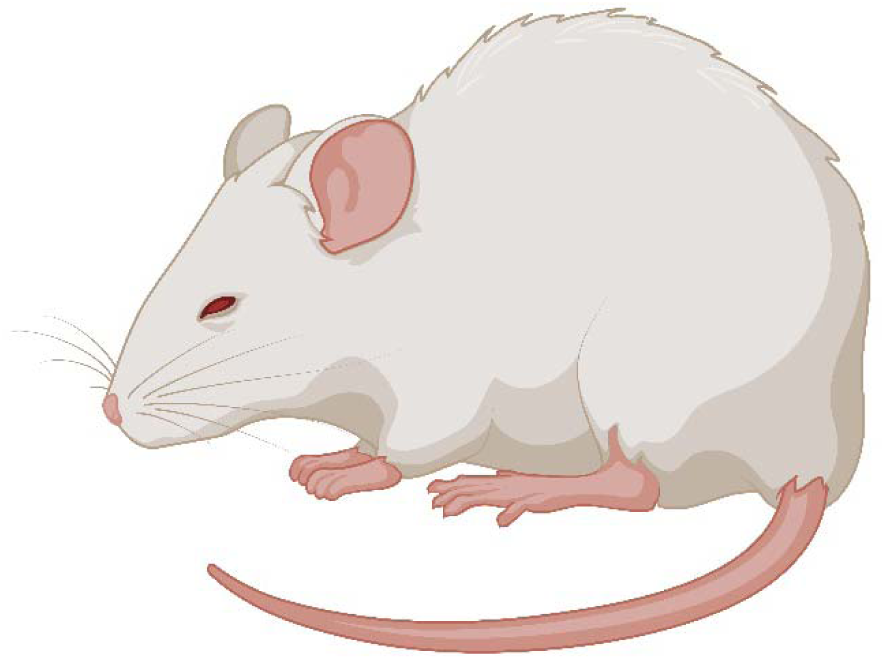
Example of sickness-associated features (ears laid back against the head, nose/cheek flattened, squinted eyes, and piloerect fur). Created with Biorender.

### 2.5 Cytokine array

Analysis of 24-hour data identified 8 hours post-LPS as the time point when the largest differences in fever and sickness behavior were observed; therefore, a separate group of rats was decapitated at eight hours post-LPS. Terminal blood was collected in EDTA tubes and centrifuged (4°C, 3000g, 15 minutes) to collect plasma. A commercially available, slide-based cytokine microarray (Abcam ab197484) was used to quantify levels of the following cytokines: IFN-γ, IL-1α, IL-1β, and IL-10. Slides were visualized using a Cy3 laser scanner (GenePix® 4000B), and median signal values from four replicates per cytokine were compared to standards.

### 2.6 Microglia reconstructions

Microglial morphology was used as a metric for the central neuroinflammatory response to LPS. Microglial deramification and decreased complexity correlate with elevated cytokine production, suggesting that this morphometric glial state is representative of a proinflammatory microglia phenotype (Althammer et al., 2020). At eight hours post-LPS, rats were decapitated, brains were removed and drop-fixed in 4% paraformaldehyde for 24 hours, followed by 30% sucrose until sectioning (Harder et al., 2023).

Fixed tissue was sectioned in a 1:6 series of 40-μm coronal sections with a Leica SM2010R microtome and stored in cryoprotectant at –20°C. Microglia were visualized using immunohistochemistry as previously described (Doyle and Murphy, 2017; Eidson and Murphy, 2013). Briefly, free-floating sections were rinsed thoroughly in potassium phosphate buffer solution (KPBS), incubated in 3% hydrogen peroxide at room temperature for 30 minutes, and then rinsed in KPBS. Sections were then incubated in 1:10,000 rabbit anti-Iba1 (Wako; 019- 19741) diluted in KPBS with 1% Triton-X overnight at room temperature. Following rinses in KPBS, sections were incubated in 1:600 biotinylated donkey anti-rabbit (Jackson Immuno; 711- 065-152) diluted in KPBS with 0.4% Triton-X for one hour at room temperature. Following KPBS rinses, sections were incubated in an Avidin/Biotin solution (PK-6100, Vector Labs) for one hour at room temperature, followed by KPBS and sodium acetate rinses. The sections were then incubated in a 3,3’-diaminobenzidine solution for 30 minutes, rinsed with sodium acetate and KPBS, and mounted onto slides. Slides were dehydrated using increasing concentrations of ethanol and cover-slipped. Microglial morphology was imaged in the ventrolateral PAG (level 4; 8.04 mm posterior to bregma; Figure 3A; blue box), given its central role in the regulation of opioid-induced immunosuppression. As an anatomical control, microglial morphology was also analyzed in the entorhinal cortex (8.04 mm posterior to bregma; Figure 3A; red box). Like the PAG, the entorhinal cortex has dense µ-opioid receptor expression, but does not have a defined role in immune signaling. One to three sections per rat were imaged unilaterally at 40x on the Keyence BZ-X700 using the Z-stacking feature (1 µm steps). Microglia were then reconstructed using Imaris 10.0.0. Images were first converted into Z-stack TIFF files in FIJI, then converted to .ims files and opened in Imaris for preprocessing (inversion and background subtraction).

**Figure 3.**
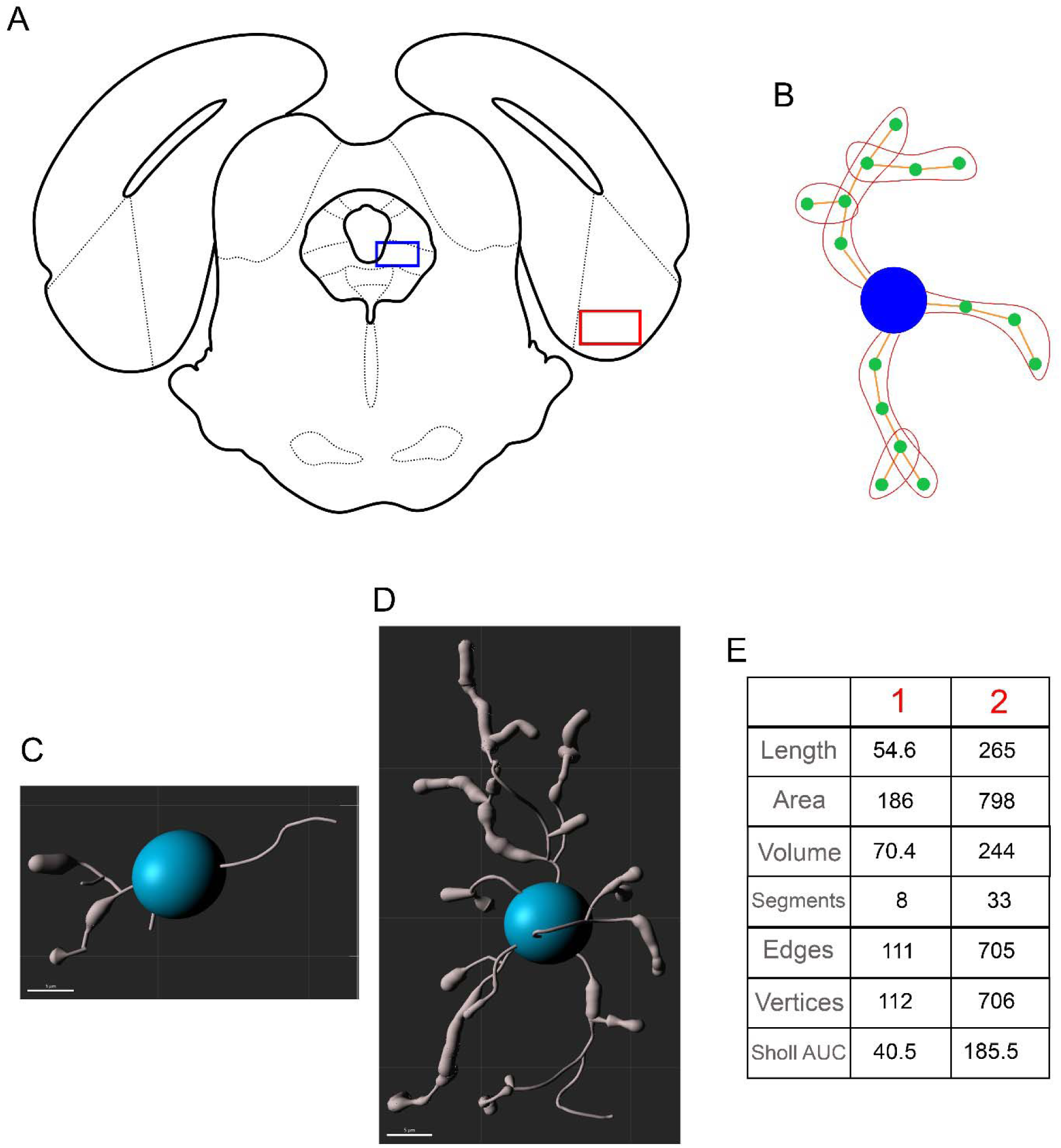
Location of vlPAG (**A, red box**) and entorhinal cortex (**A, blue box**) for analysis of microglial morphology. dlPAG = dorsolateral periaqueductal gray, DR = dorsal raphe, Ent = entorhinal cortex, FMJ = forceps major, IC = inferior colliculus, LFP = longitudinal fasciculus of the pons, lPAG = lateral periaqueductal gray, MnR = median raphe, V1 = primary visual cortex, vlPAG = ventrolateral periaqueductal gray. **B.** Example of segments, edges, and vertices. The blue circle represents the soma, orange lines represent edges, green circles represent vertices, and red outlines represent segments. Example of deramified (**C**) vs. ramified (**D**) microglia and their respective morphological values (**E**).

Microglia morphology was then traced and analyzed using the filament creation wizard, followed by manual validation. A total of 1588 microglia were reconstructed and analyzed in the PAG (VEH M: 372 microglia from 6 rats; VEH F: 296 microglia from 4 rats; MOR M: 496 microglia from 8 rats; MOR F: 424 microglia from 7 rats), and a total of 266 microglia were reconstructed and analyzed in the entorhinal cortex (VEH M: 61 microglia, VEH F: 59 microglia, MOR M: 76 microglia, VEH F: 70 microglia).

Seven metrics were collected:

1. Length: total length of all edges in the microglia (µm).
2. Area: total area of each microglia (µm).
3. Volume: total volume of each microglia (µm).
4. Segments: total number of segments in the microglia.
5. Edges: total number of connections between vertices.
6. Vertices: total number of points connecting edges.
7. Sholl intersections: number of intersections on concentric spheres spaced 1 µm apart. See Figure 3B for an example of segments, edges, and vertices. Figure 3 also shows an example of two microglia: one with deramified morphology (C) and one with ramified morphology (D), along with values of all seven metrics collected for each microglia (E). Deramified morphology is associated with smaller values, while ramified morphology leads to greater values on all seven metrics. Deramification correlates with cytokine production, suggesting that deramified microglia are in a functionally “active” state (Althammer et al., 2020). Sholl intersections were analyzed using area under the curve (AUC) such that smaller AUC values are representative of fewer Sholl intersections.

### 2.7 Gut permeability

Gut permeability was measured using oral administration of fluorescein-isothiocyanate-labeled dextran (FITC-dextran; molecular weight 4kDa; Sigma-Aldrich 46944). Adult male and female rats (P60) were fasted for four hours (beginning at nine AM). Following the fast, rats were orally dosed with 600 mg/kg of a 125 mg/mL solution of FITC-dextran using a 16g three-inch curved gavage needle with a 3mm ball. Four hours post-FITC-dextran, blood was collected from the saphenous vein into EDTA tubes and centrifuged (4°C, 3000g, 15 minutes) to separate plasma. Samples (100 µL) were transferred to a black-bottom 92 well plate, and relative fluorescent units (RFU) were read using a SpectraMax M2 plate reader (emission 530 nm, excitation 485 nm). Data were normalized within sexes to generate fold change of MOR vs. VEH rats.

### 2.8 Measurement of bacterial contact via anti-LPS antibody levels

Anti-LPS antibody levels were determined in adult male and female rats (P60) using ELISA. Blood was collected from the saphenous vein into uncoated microcentrifuge tubes, allowed to clot for 30-60 minutes, then centrifuged (4°C, 3000g, 15 minutes) to generate serum. Plates were coated in-house using a 0.5% v/w LPS and carbonate-bicarbonate buffer solution (100 µL per well) and washed the following day using 0.05% goat serum and 0.01% TWEEN20 in PBS solution. The plate was then incubated at 37°C for one hour in the presence of 100 µL serum (per well; diluted 1:200). Following a wash, 100 µL of 0.1% v/v HRP-conjugated anti-rat IgG antibody was added to each well, incubated at 37°C for one hour and then washed again. The reaction product was visualized by adding 100 µL of SureBlue TMB (SeraCare; 5120-0075) to each well. After a five minute incubation in the dark at room temperature, 100 µL of TMB stop solution (SeraCare; 5150-0020) was added. Optical density was read using a Bio-rad iMark microplate reader at 450 nm.

### 2.9 Measurement of antibody production

To investigate whether any potential differences in anti-LPS antibodies were generalized or specific, levels of the three major subtypes of antibodies (IgG, IgA, and IgM) were quantified in adult male and female rats (P60) using ELISA. Blood was collected from the saphenous vein into EDTA tubes and centrifuged (4°C, 3000g, 15 minutes) to separate plasma. Total levels of IgG (ThermoFisher; 88-50490), IgA (ThermoFisher; 88-50480), and IgM (ThermoFisher; 88-50540) were assayed following the manufacturer’s protocols. Optical density was read using a Bio-rad iMark microplate reader at 450 nm and compared to standard curves to calculate IgG, IgA, and IgM concentrations in ng/mL.

### 2.10 Experimental design and statistical analysis

Significant effects of sex, treatment, and time (where applicable) were assessed using two- or three-way mixed models or repeated measures mixed models; p<0.05 was considered significant. As repeated measures ANOVA cannot handle missing values, data were analyzed by fitting mixed models with Greenhouse-Geisser correction as implemented in GraphPad Prism 9.1.0 (Motulsky, 2023). Tukey’s or Sidak’s post-hoc tests were conducted to determine significant mean differences between *a priori* specified groups. Due to the method of partitioning variance in linear mixed models, there is no universal method to calculate standardized effect sizes (e.g., η2 for ANOVA). Whenever possible, we report unstandardized effect sizes, which agrees with recommendations for effect size reporting (Pek and Flora, 2018), including the guidance of the American Psychological Association Task Force on Statistical Inference (Wilkinson, 1999).

As multiple microglia were reconstructed and analyzed from one rat, the assumption of independence was not met, and traditional statistical analyses could not be utilized. Therefore, hierarchical bootstrapping and permutation testing were used, which do not require independent observations (see Saravanan et al., 2020 for more details on hierarchical data in neuroscience and the application of bootstrapping for this type of data). Analyses were completed in R 4.2.3 using the package ClusterBootstrap (Deen and de Rooij, 2020). General linear models were utilized to create estimated means, and permutation testing was completed to investigate the robustness of the estimated cluster means. Data were visualized as a probability density function implemented in Excel 2019. Leftward shifts in population density function are indicative of increased activation in response to LPS.

All unstandardized effect sizes are reported as differences between means (MOR-VEH) ± standard error of the mean. Female rats used to generate offspring were randomly assigned to the MOR or VEH condition. All experiments included both male and female offspring. No differences were observed between rats of different litters in the same drug exposure group (i.e., MOR and VEH); therefore, individual rats from the same litter across multiple litters served as a single cohort (4 VEH litters, 5 MOR litters). All analyses were completed blinded to the treatment group.

## 3. Results

### 3.1 24-hour LPS-induced fever

We first investigated the impact of perinatal morphine exposure on the response to LPS, focusing initially on fever and sickness behavior. After the prototypical spike in temperature due to handling stress (Machado et al., 2020), systemic administration of LPS induced a febrile response in both MOR and VEH groups (Figure 4A). Body temperature began to rise for both treatment groups at hour 2 and continued to increase from hours 3-6, with MOR males displaying a slower rise in temperature. Beginning at hour 6, body temperature began to decline for VEH males and females; in contrast, MOR males and females continued to increase and remained elevated throughout hours 7-8 before returning to baseline. By hour 11, body temperature was comparable for all rats, and by hour 16 all rats returned to baseline.

**Figure 4.**
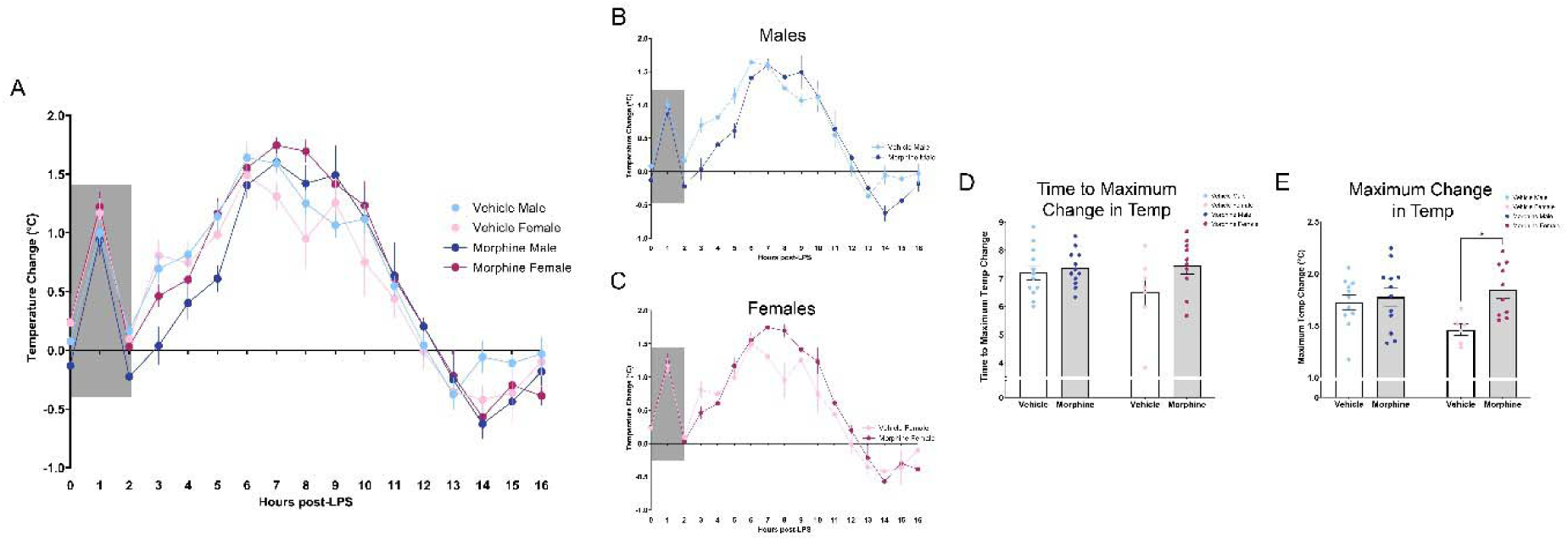
Perinatal morphine exposure leads to increased fever response to LPS. **A.** Fever arc from hours 0-16 post-LPS. **B.** MOR males showed a lower rise in body temperature from hours 3-6; temperature remained elevated vs. VEH males from hours 7-9. **C.** MOR females have elevated temperatures from hours 7-12. **D.** MOR males and females have delayed time to reach maximum fever. **E.** MOR males and females have increased maximum fever magnitude. N_VEH M_ = 12, N_VEH F_ = 7, N_MOR M_ = 12, N_MOR F_ = 10. * = significant at p<0.05.

Overall, we observed a significant effect of treatment on LPS-induced fever [Time*Txt, F(24,520)=2.290, p=0.0005]; this difference was primarily driven by females [Txt*Sex, F(1,37)=10.81, p=0.0022], with MOR females maintaining a higher fever response for a longer period of time in comparison to VEH females (Figures 4B-C). We next analyzed two components of the fever response: time to maximum change in temperature and maximum change in temperature. For time to maximum change in temperature, although we observed no significant effect of treatment [Txt, F(1,37)=3.508, p=0.0690], MOR females took longer to reach maximum fever (Figure 4D). Specifically, MOR female rats took 0.95 hours longer (VEH F vs MOR F, p=0.1811), shifting from an average time of 6.5 hours to 7.45 hours. No differences were noted for males (VEH M vs MOR M, p=0.9722). A significant main effect of treatment was observed in maximum change in temperature [Txt, F(1,35)=6.537, p=0.0151] (Figure 4E); this effect was also driven by females (VEH M vs. MOR M, p=0.9604; VEH F vs. MOR F, p=0.0318). The maximum temperature change for MOR F was 1.84°C vs. 1.46°C for VEH F, a 0.38°C difference, equivalent to a 26.2% greater temperature change.

Together, this data suggests that perinatal opioid exposure leads to increased febrile response to LPS. MOR males and females take longer to reach maximum fever and have a higher maximum fever, both of which were only significant in females.

### 3.2 24-hour sickness behavior

Similar to humans, rodents also display sickness-associated features in response to infections that function to conserve energy and aid in recovery and survival. To characterize this quantitatively, we utilized a Likert scale based on the presence/absence and severity of four physical attributes associated with sickness (Sotocinal et al., 2011). LPS induced robust sickness in VEH male rats and MOR male and female rats (Figure 5A); in contrast, VEH female rats showed minimal signs of sickness despite robust fever generation post-LPS [Time*Txt*Sex, F(11,165)=2.170, p=0.0183]. Overall, MOR females showed higher sickness scores vs. VEH females from hours 4-14; no significant differences were noted in MOR vs. VEH males (Figures 5B-C). MOR females also showed higher maximum sickness scores vs. VEH females [Txt, F(1,15)=12.36, p=0.0031; Sex, F(1,15)=5.815, p=0.0292], while males in both treatment groups showed comparable levels of sickness (Figure 5D). Importantly, sickness scores were positively correlated with maximum temperature change [r(41)=0.4268, p=0.0043], confirming that our sickness scale captured the relevant features associated with LPS-induced fever (Figure 5E).

**Figure 5.**
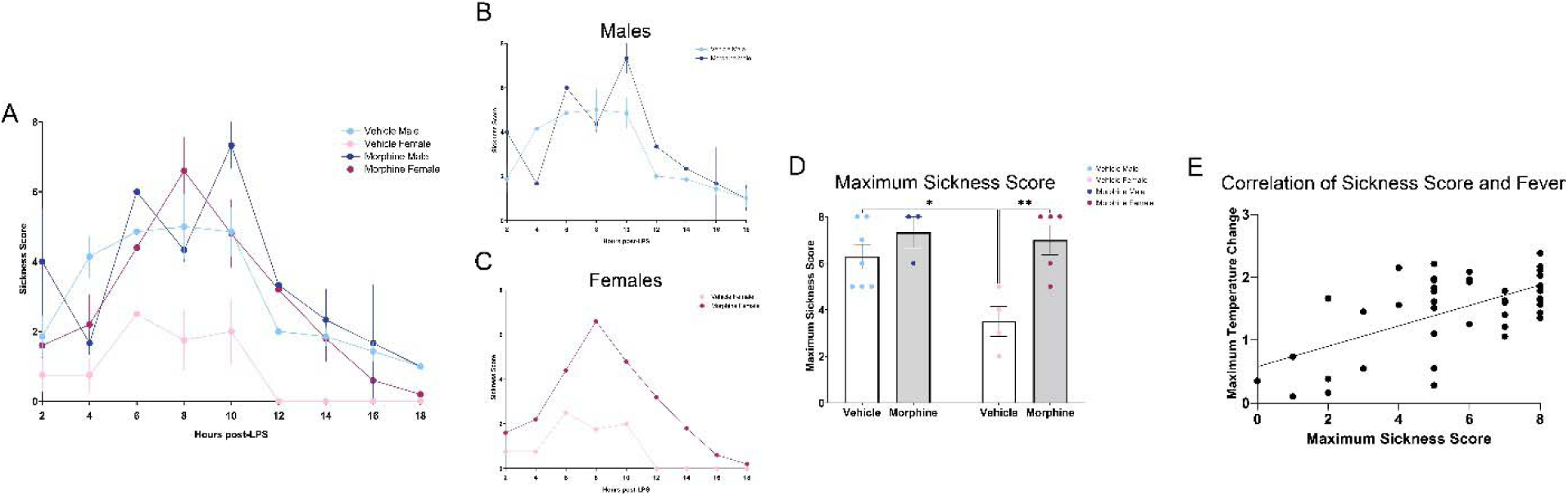
Perinatal morphine exposure leads to elevated sickness behavior in female rats. **A.** Sickness arc from hours 0-16. **B.** No difference was observed in males. **C.** MOR females have elevated sickness behavior. **D.** MOR females have elevated maximum sickness behavior. **E.** Maximum temperature and maximum sickness scores positively correlate. N_VEH M_ = 9, N_VEH F_ = 7, N_MOR M_ = 12, N_MOR F_ = 10. * = significant at p<0.05.

Together, this data suggests that LPS-induced fever and sickness is increased in MOR male and female rats. We next examined potential mechanisms by which perinatal opioid exposure leads to an increased response to LPS.

Based on the fever arc for VEH and MOR rats, we chose the 8 hour timepoint for analyses of central and peripheral immune measures. At this timepoint, both male and female MOR rats had significantly elevated body temperature [Txt, F(1,37)=8.844, p=0.0052], although only female rats reached statistical significance (VEH M vs. MOR M, p=0.8196; VEH F vs. MOR F, p=0.0167) (Figure 6A). Body temperature at 8 hours post-LPS was 13.7% higher in MOR males (MOR M: 1.42±0.16°C; VEH M: 1.25±0.11°C) and 78.4% higher in MOR females (MOR F: 1.70±0.096°C; VEH F: 0.95±0.26°C). Sickness scores were similarly elevated, again only in MOR females [Txt*Sex, F(1,39)=7.69, p=0.0088] (Figure 6B). Specifically, the average sickness score for VEH females was 1.7; in contrast, the average sickness score for MOR females was 4.5, over 2.5 times greater [p=0.0325].

**Figure 6.**
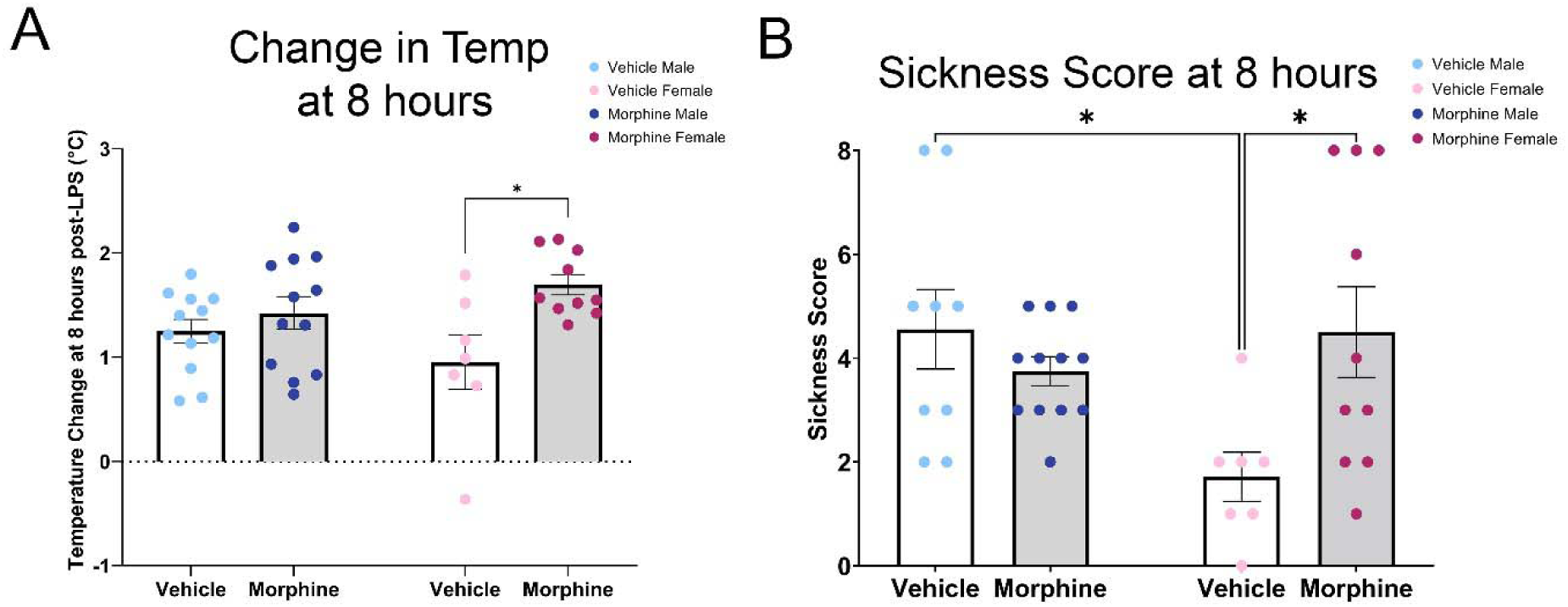
Perinatal morphine exposure leads to increased fever and sickness scores 8 hours post-LPS. **A.** MOR males and females have elevated body temperature at 8 hours post-LPS. **B.** MOR females have elevated sickness behavior at 8 hours post-LPS. N_VEH M_ = 9-12, N_VEH F_ = 7, N_MOR M_ = 12, N_MOR F_ = 10. * = significant at p<0.05.

### 3.3 Cytokine array

We first tested the hypothesis that perinatal exposure to morphine leads to long-term immune dysregulation, focusing initially on plasma cytokine levels, which constitute a major component of an organism’s response to LPS and infection, and are responsible for fever induction. Our analysis focused on four cytokines implicated in inflammation and response to infection: IFN-γ, IL-1α, IL-1β, and IL-10. A significant effect of treatment was only observed in one cytokine, IL-1α [Txt, F(1,18)=10.96, p=0.0039] (Figure 7A), with concentrations three times greater in MOR vs. VEH rats. IL-1α acts primarily in the hypothalamus to generate fever, suggesting that the elevated level of this cytokine may be the primary driver of the elevated fever observed in MOR rats. Given the variability in the levels of IL-1α in MOR rats, we next investigated if IL-1α levels were correlated with fever response. However, IL-1α was not correlated to temperature change 8 hours post-LPS (Figure 7B; r(22)=0.3032, p=0.1701). We also investigated levels of IFN-γ, IL-1β, and IL-10; however, no significant treatment differences were observed (Figure 7C-E).

**Figure 7.**
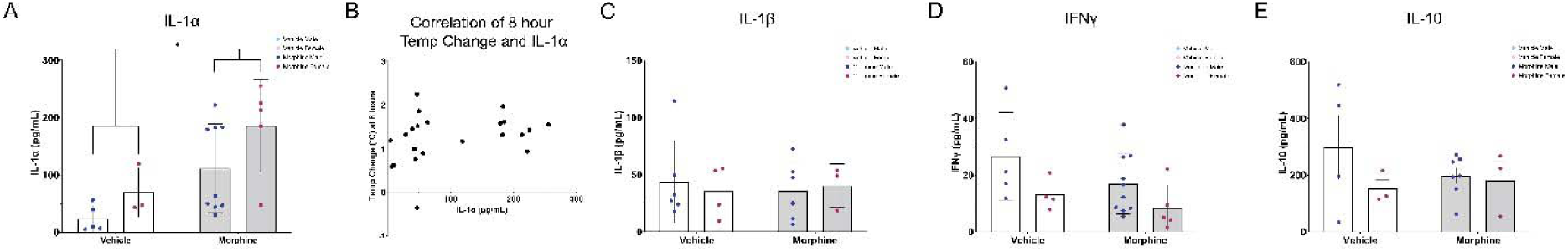
Alterations in cytokine levels post-LPS. **A.** Elevated IL-1α levels in MOR rats. **B.** No correlation of IL-1α with temperature change 8 hours post-LPS. **C.** No changes in IL-1β. **D.** No changes in IFNγ. **E.** No changes in IL-10. N_VEH M_ = 4-6, N_VEH F_ = 3-4, N_MOR M_ = 6-10, N_MOR F_ = 3-5. * = significant at p<0.05.

### 3.4 Microglia reconstructions

Although morphine is typically considered immunosuppressive in the periphery (decreased natural killer cell cytotoxicity, reduced macrophage phagocytosis, and altered proinflammatory cytokine production), centrally, morphine promotes microglial reactivity and initiates cytokine release in a TLR4-dependent manner (Eidson et al., 2017; Wang et al., 2012). Thus, we next examined if perinatal morphine alters the microglial response to LPS.

Microglia respond to LPS by transforming to a more “activated” and deramified morphology, reflected as smaller size and less complex structure. Thus, we predicted that microglia would show increased deramified morphology in MOR rats. Microglia morphology was reconstructed, and seven metrics were analyzed: length, area, and volume; segments, edges, and vertices; and Sholl intersections (see Figure 8 for representative microglia traces from all four groups). In all metrics, smaller measurements represent deramified and “activation-associated” morphology.

**Figure 8.**
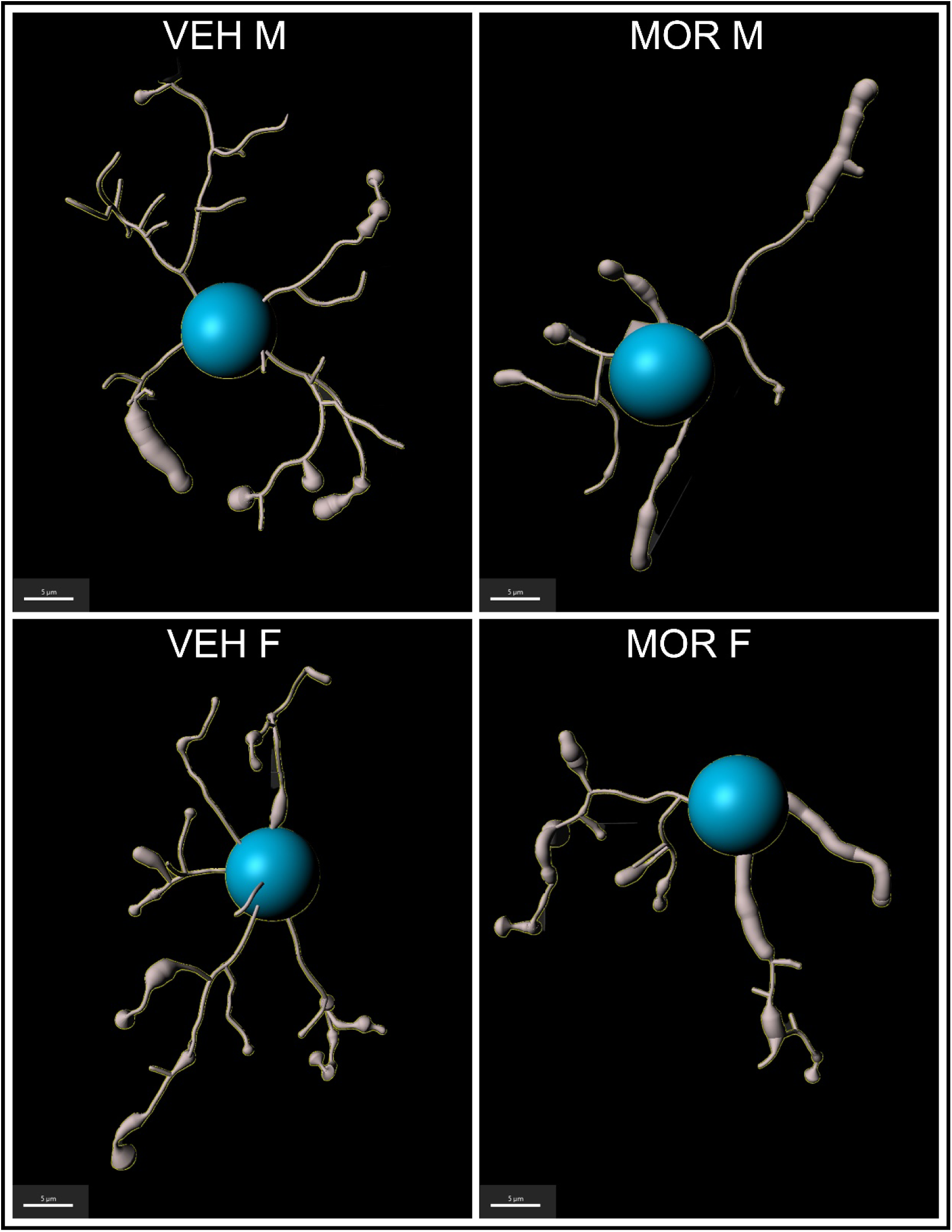
Representative traces of microglia from the PAG of VEH M, VEH F, MOR M, and MOR F.

We first analyzed microglia size (length, area, and volume) and observed a leftward population shift for all three measures in MOR rats independent of sex (Figure 9A-C).

**Figure 9.**
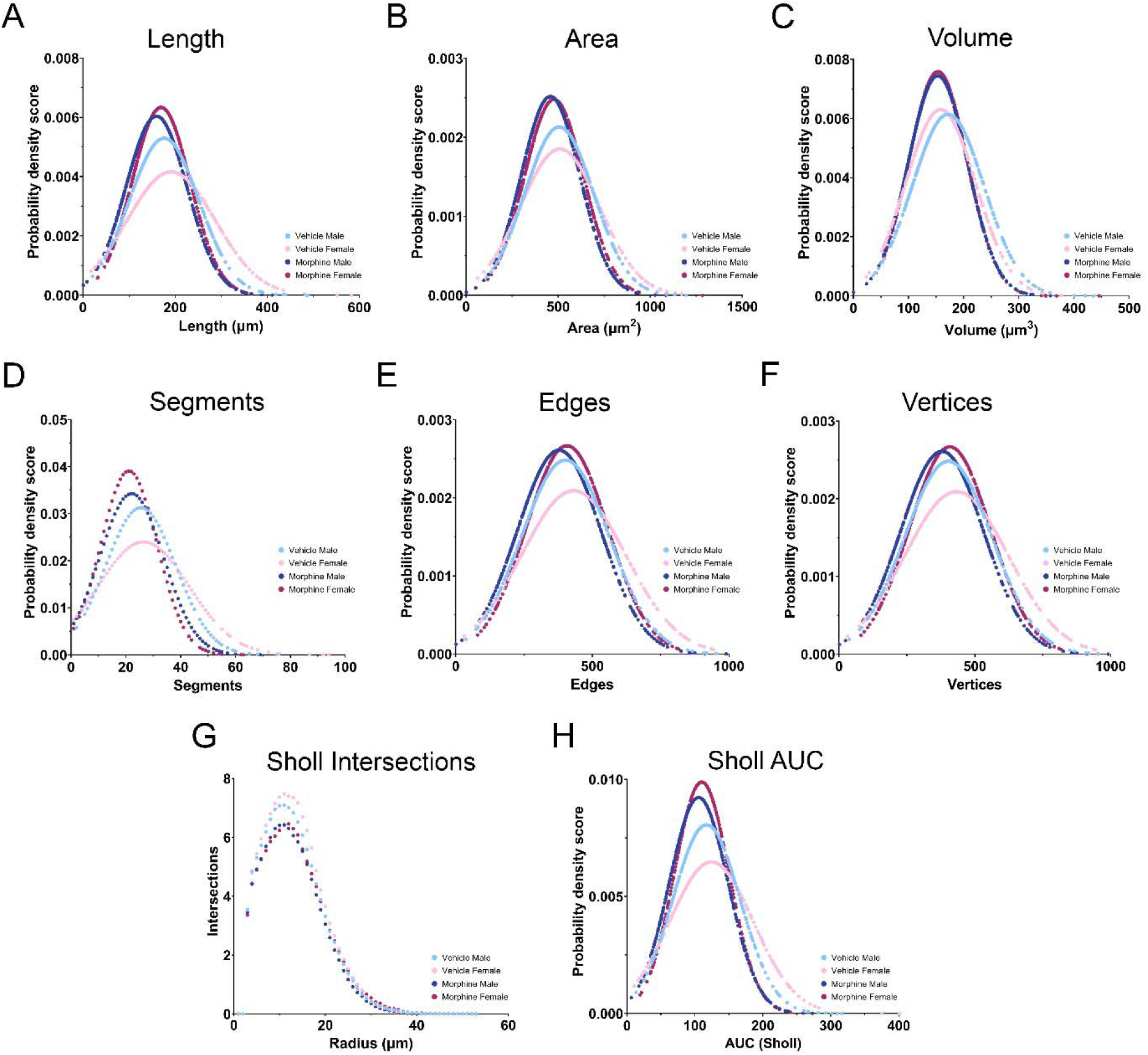
Microglia activation 8 hours post-LPS. A-C. Length, area, and volume are all decreased in MOR male and female rats. **D-F.** The number of segments, edges, and vertices is decreased in MOR male and female rats. **G.** MOR male and female rats have fewer intersections across the entire Sholl radius. **H.** Sholl AUC is decreased in MOR male and female rats. N_VEH M_ = 372 microglia, N_VEH F_ = 296 microglia, N_MOR M_ = 496 microglia, N_MOR F_ = 424 microglia.

Hierarchical bootstrapping followed by permutation testing identified a significant difference in mean microglial length of -17.04 µm (-9.6%) and -18.82 µm (-10%) for MOR male and female rats, respectively, in comparison to VEH rats. Similar results were observed for both area and volume: for area, mean differences of -49.64 µm^2^ (-9.8%) and -25.4 µm (-5.1%) were observed in MOR male and female rats; for volume, MOR male and female rats had mean differences of - 19.33 µm^3^ (-11.2%) and -3.1 µm (-2.0%). See Table 1 for descriptive statistics for all metrics analyzed. Overall, the observed leftward shift in microglia size for MOR rats suggests that POE increases microglial activation-associated morphology (i.e., deramification) in response to LPS.

**Table 1.**
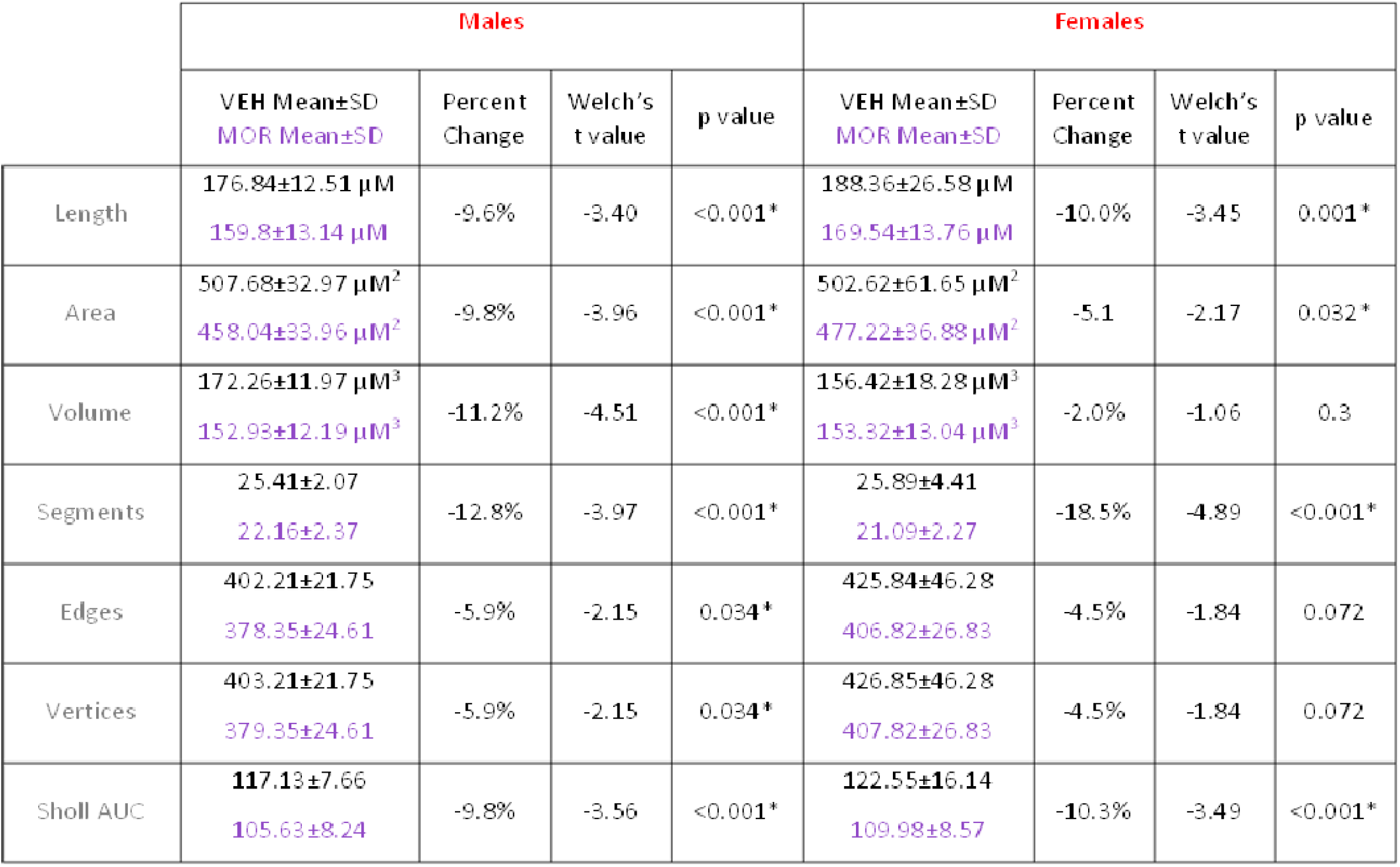
Summary of alterations in microglial morphology in the periaqueductal gray. % represent percent change from sex-matched VEH controls. * = significant at p<0.05.

We also analyzed additional measures of size and complexity, including the number of segments, edges, and vertices present per microglia (Figure 9D-F). Consistent with the results observed for size, all measures were reduced in MOR vs. VEH rats. Mean differences of -3.25 (- 12.8%) and -4.8 segments (-18.5%) were observed in MOR male and female microglia. For edges, mean differences of -23.86 (-5.9%) and -19.02 (-4.5%) were noted for MOR vs. VEH rats, and for vertices mean differences of -23.86 (-5.9%) and -19.03 (-4.5%) were observed. Overall, MOR male and female rats again showed left-shifted populations of microglia size and complexity, consistent with increased activation.

We next analyzed Sholl intersections, a classic metric of microglial activation. Comparison of the number of intersections per concentric circle (1 µm apart) showed lower intersections across the entire range for MOR male and female microglia, consistent with deramification (Figure 9G). To analyze Sholl intersections via hierarchical bootstrapping, we utilized area under the curve (AUC) of individual microglia. Overall, MOR male and female microglia had lower Sholl AUC values (Figure 9H), representing fewer intersections and suggesting increased activation.

Together, all seven metrics of activation-associated morphology in microglia were significantly lower for MOR male and female rats vs. VEH controls, suggesting that perinatal opioid exposure potentiated the microglial response to LPS.

To determine whether the increased microglial activation observed in the PAG of MOR rats was specific to the PAG, we next examined microglial morphology in the entorhinal cortex, a region with high µ-opioid receptor expression but no known role in immunity. Surprisingly, results for the entorhinal cortex were similar to what was observed in the PAG: in all seven metrics, MOR male and female rats display increased activation-associated microglial morphology (see Supplemental Table 1 for a summary of these results). This suggests that increased microglial activation in response to LPS may be a generalized response in brain regions with high µ-opioid receptor expression (PAG and entorhinal cortex) for male and female rats perinatally exposed to morphine.

Thus far, we have reported that perinatal opioid exposure potentiates the response to LPS, as indicated by increased fever and sickness, elevated levels of IL-1α and increased microglial activation. We next examined if these differences were related to basal differences in immune function, such that MOR rats are less able to launch an appropriate immune response following exposure to antigens or pathogens. We first investigated gut permeability using FITC-dextran dissemination into the bloodstream. Bacteria in the gut are a major source of immune system stimulation and training, and increased gut permeability would promote increased bacterial contact with the immune cells in the lamina propria, leading to differential immune system development (Kaczmarczyk et al., 2021). In addition, opioids are known to slow gut peristalsis, which is associated with increased gut permeability (Akbarali and Dewey, 2019).

### 3.5 Analysis of gut permeability using FITC-dextran

Due to differences in basal levels between experimental rounds, data were normalized to the mean of the VEH group to generate fold changes for each sex. Overall, no significant differences in gut permeability were noted (Figure 10A-B). However, MOR females had an average fold change of 1.47 relative to VEH females, representing a 47.1% increase in gut permeability [unpaired T test; t(16)=2.038, p=0.0585]; no differences were observed for males [fold change 0.79; unpaired T test; t(14)=1.505, p=0.1546]. This suggests that exposure to morphine *in utero* leads to long-term increases in gut permeability for female rats, which may alter immune system development and impact the response to immune stimulators.

**Figure 10.**
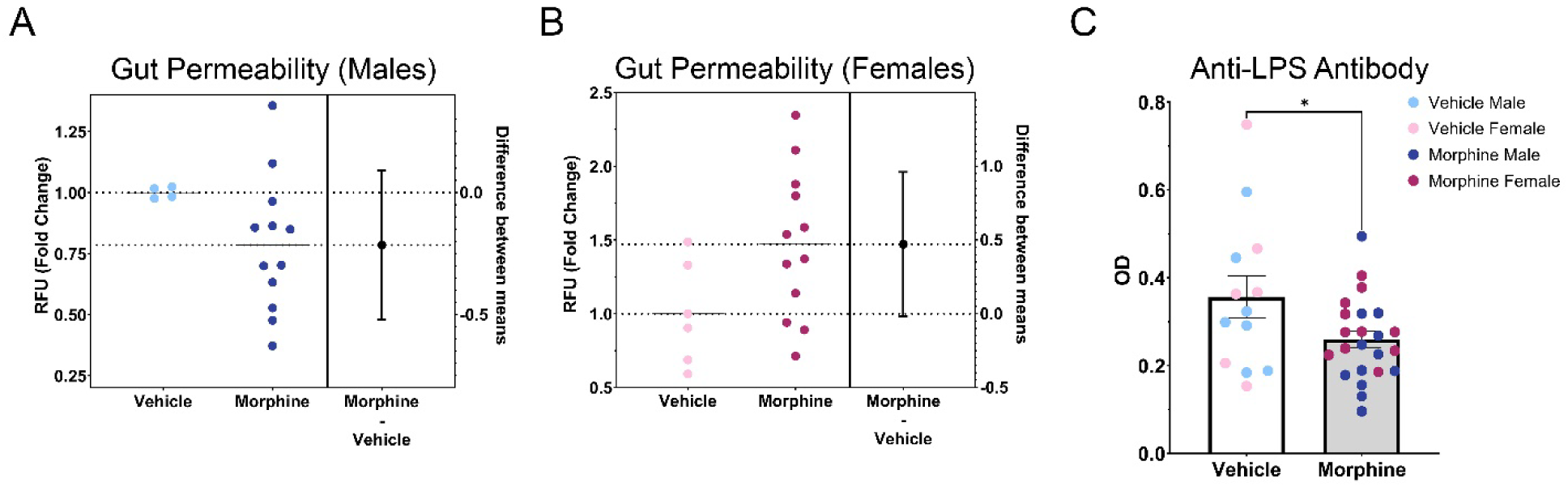
MOR female rats have increased gut permeability; however, both male and female MOR rats have decreased anti-LPS antibody levels. **A.** No differences in gut permeability were found between males. **B.** MOR females have elevated gut permeability. **C.** MOR males and females have decreased anti-LPS antibody levels, despite elevated gut permeability seen in females. N_VEH M_ =4-5, N_VEH F_ = 5-6, N_MOR M_ = 8-12, N_MOR F_ = 10-12. * = significant at p<0.05.

### 3.6 Measurement of bacterial contact via anti-LPS antibody levels

To confirm our observed, albeit non-significant, increases in gut permeability in MOR female rats, we next quantified levels of anti-LPS antibodies. Increased gut permeability would allow for increased bacterial dissemination into the lamina propria; as LPS is one of the major antigens utilized by the immune system to recognize bacteria, we predicted that levels of anti-LPS antibodies would be similarly elevated in MOR females. We observed no significant effect of sex, so males and females were combined to increase power. In contrast to our predicted increase, we observed a significant decrease in anti-LPS antibodies in MOR rats [t(34)=2.191, p=0.0354] (Figure 10C). Anti-LPS antibody levels were, on average, 27.1% lower in MOR vs. VEH rats. As the increased gut permeability observed in MOR females is in conflict with the observed decrease in anti-LPS antibodies, we next measured levels of antibody classes IgG, IgA, and IgM to identify potential deficits in antibody production as an alternative explanation for decreased anti-LPS antibody levels.

### 3.7 Measurement of antibody production

We first analyzed IgG, the most common and abundant antibody subtype, and a primary mechanism to target microbes for phagocytosis. Our analysis identified a significant interaction of treatment and sex [F(1,25)=9.525, p=0.0049] (Figure 11A). Specifically, we noted a 56% reduction in IgG production in MOR females (VEH F: 7.20*10 ±1.84*10 ng/mL; MOR F: 3.14*10 ±6.08*10 ng/mL; p= 0.0002). Similar mean differences were observed in MOR males, with a reduction of 30% (VEH M: 2.50*10 ±6.12*10 ng/mL; MOR M: 1.75*10 ±2.31*10 ng/mL; p=0.6950). We also report that IgG levels were three times higher in VEH females vs. VEH males (p<0.0001), consistent with results showing that females typically have a more robust antibody response and higher baseline immunoglobulin levels (Klein and Flanagan, 2016).

**Figure 11.**
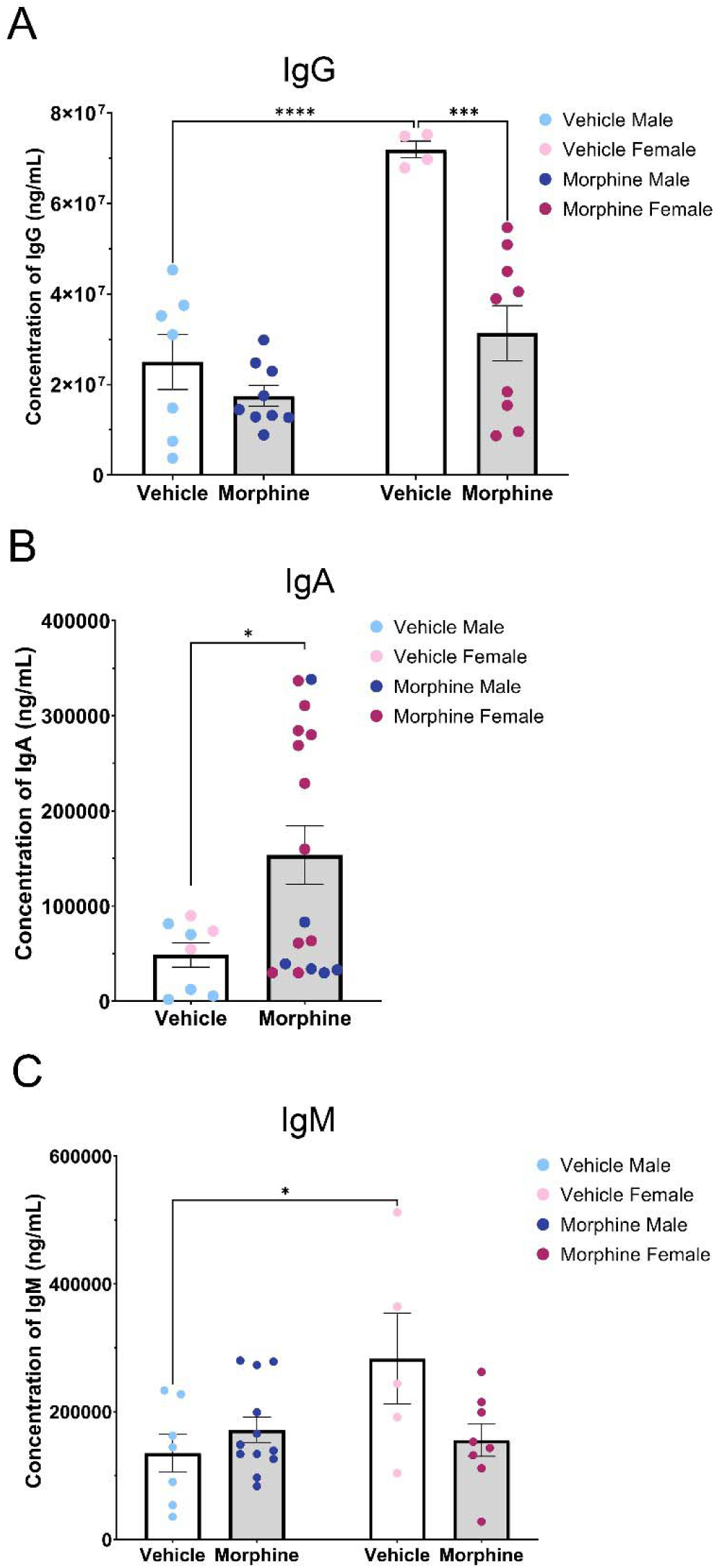
Perinatal opioid exposure alters antibody levels. **A.** MOR females have decreased levels of IgG. **B.** MOR male and female rats have elevated IgA antibody levels. **C.** MOR female rats trend toward lower levels of IgM. N_VEH M_ = 5-7, N_VEH F_ = 3-5, N_MOR M_ = 6-12, N_MOR F_ = 8-11. * = significant at p<0.05.

We next analyzed IgA, which is involved in mucosal immunity and defense against oral or respiratory pathogens. There was no significant effect of sex, so males and females were collapsed to increase power. IgA levels were significantly higher in MOR rats [t(23)=2.288, p=0.0317] than VEH rats (VEH: 48669±12854 ng/mL; MOR: 153596±30551 ng/mL; 215% increase; Figure 11B).

Last, we analyzed IgM, a stimulator of the classical complement system. Two-way ANOVA identified a significant interaction of treatment and sex [F(1,28)=5.974, p=0.0211] (Figure 11C). In MOR females, IgM levels were reduced by 45.1% (VEH F: 283180±71072.20 ng/mL; MOR F: 155501.25±25274.86 ng/mL, p=0.0853), while a non-significant increase was noted in males [26.8% increase; VEH M: 135285.71±29888.72 ng/mL; MOR M: 171505±20246.52 ng/mL; p=0.8334]. We again noted that VEH females had IgM levels twice that of VEH males [p=0.0432].

Together, this data suggests that perinatal opioid exposure results in generalized deficits in antibody production, observed both in anti-LPS antibodies and IgG and IgM (females only) levels. Interestingly, a significant increase in IgA levels was noted for MOR rats, potentially related to alterations in enteric immunity and gut permeability.

Results from measurements of baseline immune functioning in MOR vs. VEH rats suggest that *in utero* exposure to morphine leads to long-term alterations in immune system activity, including increased gut permeability and decreased antibody production. These differences may explain the increased fever and sickness responses to LPS and altered cytokine levels and microglial activation post-LPS. See Table 2 for a summary of the results.

**Table 2.** Summary of results.

|  | MOR M | MOR F |
| --- | --- | --- |
| Fever | ↑ | ↑ |
| Sickness | --- | ↑ |
| Cytokine Levels (IL-1α) | ↑ | ↑ |
| Microglia Activation | ↑ | ↑ |
| Gut Permeability | --- | ↑ |
| Anti-LPS Antibody | ↓ | ↓ |
| IgG | ↓ | ↓ |
| IgA | ↑ | ↑ |
| IgM | --- | ↓ |

## 4. Discussion

Previous clinical studies indicate that infants exposed to opioids *in utero* are at a higher risk of infection and hospitalization (Arter et al., 2021; Uebel et al., 2015; Witt et al., 2017). However, to date, the underlying mechanism by which perinatal opioid exposure leads to this increased risk is unknown. The present study was designed to address this gap, and, in particular, to characterize the physical and immunological response to LPS using a preclinical model of perinatal opioid exposure. We hypothesized that perinatal opioid exposure would compromise gut permeability and antibody production, consistent with the known immunosuppressive effects of chronic opioids in adult humans and rodents. Furthermore, we hypothesized that these effects would alter the response to an immune stimulator, LPS, observed as changes in fever, sickness, and markers (peripheral and central) of inflammation.

We first investigated the effects of perinatal opioid exposure on the response to an experimental model of Gram-negative bacterial infection, LPS. Our studies revealed that MOR rats responded with elevated fever and sickness following LPS administration vs. their VEH counterparts. Although increased fever and sickness may seem contradictory to the well-characterized immunosuppressive effects of opioids, fever is widespread among immunosuppressed patients and may even be the only symptom of infection (Pizzo, 1999). Thus, while healthy individuals can regulate their immune system properly, immunosuppressed patients, particularly those who are neutropenic (i.e., with low levels of neutrophils), frequently show rapid rises in core body temperature and quickly proceed to sepsis. Indeed, previous studies investigating the response to LPS in rats made leukopenic (i.e., with a low number of circulating leukocytes) via chemotherapy have reported increased fever response to LPS, along with alterations in cytokine production (Miñano et al., 2004; Tavares et al., 2006, 2005). Interestingly, the time course and magnitude of fever in these studies are similar to what is observed in the present study for MOR rats. This, along with our observed deficits in antibody production, suggests that perinatal opioid exposure may produce neutropenia and/or leukopenia.

In addition to the febrile response, the majority of rats that received LPS also displayed physical characteristics of sickness. Specifically, male rats and MOR female rats looked sick, with ears flattened back, eyes tightened, nose flattened, and piloerect fur. Surprisingly, female VEH rats displayed very few physical attributes of sickness, despite a robust fever response. No difference in sickness score was noted for MOR vs. VEH males, perhaps due to a potential ceiling effect, given the high sickness scores observed in VEH males. Male rodents generally exhibit more severe sickness behavior following LPS, including anorexia (Kuo, 2016; Pitychoutis et al., 2009), huddling, piloerection, ptosis, lethargy (Cai et al., 2016), and reduced locomotion (Yee and Prendergast, 2010); however see (Pitychoutis et al., 2009). For the present studies, we chose to use a low dose of LPS that would enable us to observe either an increase or decrease in sickness score. Use of even lower doses of LPS may elucidate whether our observed sex difference in VEH rats is due to increased sensitivity to the sickness-promoting effects of LPS in males.

Peripheral cytokine levels were also altered in response to LPS treatment. Elevation of IL-1α levels in MOR male and female rats at eight hours post-LPS likely contributes to the potentiated fever response seen in MOR rats, as IL-1α binds to IL-1R in the anterior and paraventricular nucleus of the hypothalamus and initiates a downstream cascade ending in synthesis of the pyrogen prostaglandin E2 (Cartmell et al., 1999), which increases body temperature via brown adipose tissue metabolism and vasoconstriction. Previous studies have indicated that acute morphine, given in conjunction with LPS, led to an increase in IL-1α levels in the brain (Roy et al., 1999), potentially through synergy with TLR4 (Eidson et al., 2017). Other models of perinatal opioid exposure have reported increases in adult levels of TLR4 and MyD88 (Jantzie et al., 2019; Smith et al., 2022). As IL-1α is one of the primary cytokines released after TLR4 stimulation via NF-κB signaling, this provides a potential mechanism by which perinatal morphine exposure increases IL-1α levels. We also noted a high degree of variability in IL-1α levels; surprisingly, this was not related to maximum temperature induced by LPS, but rather, this may be associated with the degree of immunosuppression in MOR rats, including elevations of TLR4 and/or MyD88.

The current study also identified increased microglial activation in the ventrolateral periaqueductal gray in male and female MOR rats following LPS treatment, suggesting that POE leads to long-term changes in microglial reactivity. While opioid exposure typically promotes peripheral immunosuppression, previous studies have shown that microglia are generally activated by morphine in a TLR4-dependent manner (Doyle et al., 2017). Increased microglial activation likely contributed to the increased fever and sickness behavior following LPS, as activated microglia release cytokines that act in the hypothalamus to promote fever and sickness behavior. In the present study, only peripheral cytokines were assessed. Here, we observed a significant increase in IL-1α; however, plasma cytokine levels are not always representative of local brain region concentrations, so future studies should investigate local cytokine levels in the hypothalamus and PAG. We also analyzed microglial phenotype in the entorhinal cortex to determine if the observed increase in microglial activation was specific for the PAG, or a more widespread phenomenon. Surprisingly, microglia in the entorhinal cortex of both male and female rats showed increased activation-associated morphology. The entorhinal cortex is primarily associated with memory formation and learning (Maass et al., 2015) and to date has not been implicated in immune signaling. As both the PAG and entorhinal cortex have high levels of µ-opioid receptor expression, this suggests that perinatal morphine may act on µ- opioid receptors in multiple regions of the CNS to decrease the threshold for microglial reactivity, potentially through morphine’s action as a developmental stressor (Carloni et al., 2021). Future studies should investigate microglial reactivity in additional brain regions, both with and without µ-opioid receptor expression.

Given the observed changes in LPS-induced fever and sickness in MOR rats, we next investigated if these changes were due to alterations in basal peripheral immune function or specifically a result of immune stimulation. We hypothesized that morphine exposure would promote gut permeability and increase the level of anti-LPS antibodies due to increased bacterial dissemination into circulation and that these together would potentiate the response to LPS. Here, we report that MOR rats displayed increased gut permeability, but surprisingly produced fewer anti-LPS antibodies. The reduced level of antibodies was not specific to anti-LPS, as MOR rats had lower levels of the antibody subtypes IgG and IgM (females only), suggesting an overall deficit in antibody production that would predispose these rats to infection. Interestingly, we also observed significantly increased IgA antibody levels in MOR male and female rats. This may be related to morphine’s effects on the gut, including gut permeability, as IgA is primarily involved in mucosal immunity and response to oral pathogens. Together, our observed deficits in IgG and IgM are consistent with clinical data reporting increased hospitalization rates for infection in children born with *in utero* opioid exposure (Arter et al., 2021; Uebel et al., 2015; Witt et al., 2017). The cause of antibody production deficits, including a decrease in the number of antigen-presenting cells and/or B lymphocytes involved in antibody production or the ability of these cells to produce antibodies in response to antigen stimulation, warrants further investigation. As increased gut permeability is also associated with changes in gut microbiota composition, the impact of perinatal morphine exposure on gut microbiota composition and its relationship to other immune parameters should also be examined.

Overall, our data suggests that rats perinatally exposed to morphine have an immunosuppressed phenotype (specifically increased gut permeability and deficits in antibody production) that may increase the susceptibility of the immune system to a pathogen or immune challenge, consistent with our finding that MOR rats show a potentiated fever and sickness response to LPS (see Figure 12 for a summary of the results). These results provide further evidence that exposure to opioids *in utero* leads to long-term immune deficits. As the number of infants born to mothers using opioids during pregnancy continues to rise, determining the underlying mechanism whereby these infants are more vulnerable to potential pathogen exposure is critical.

**Figure 12.**
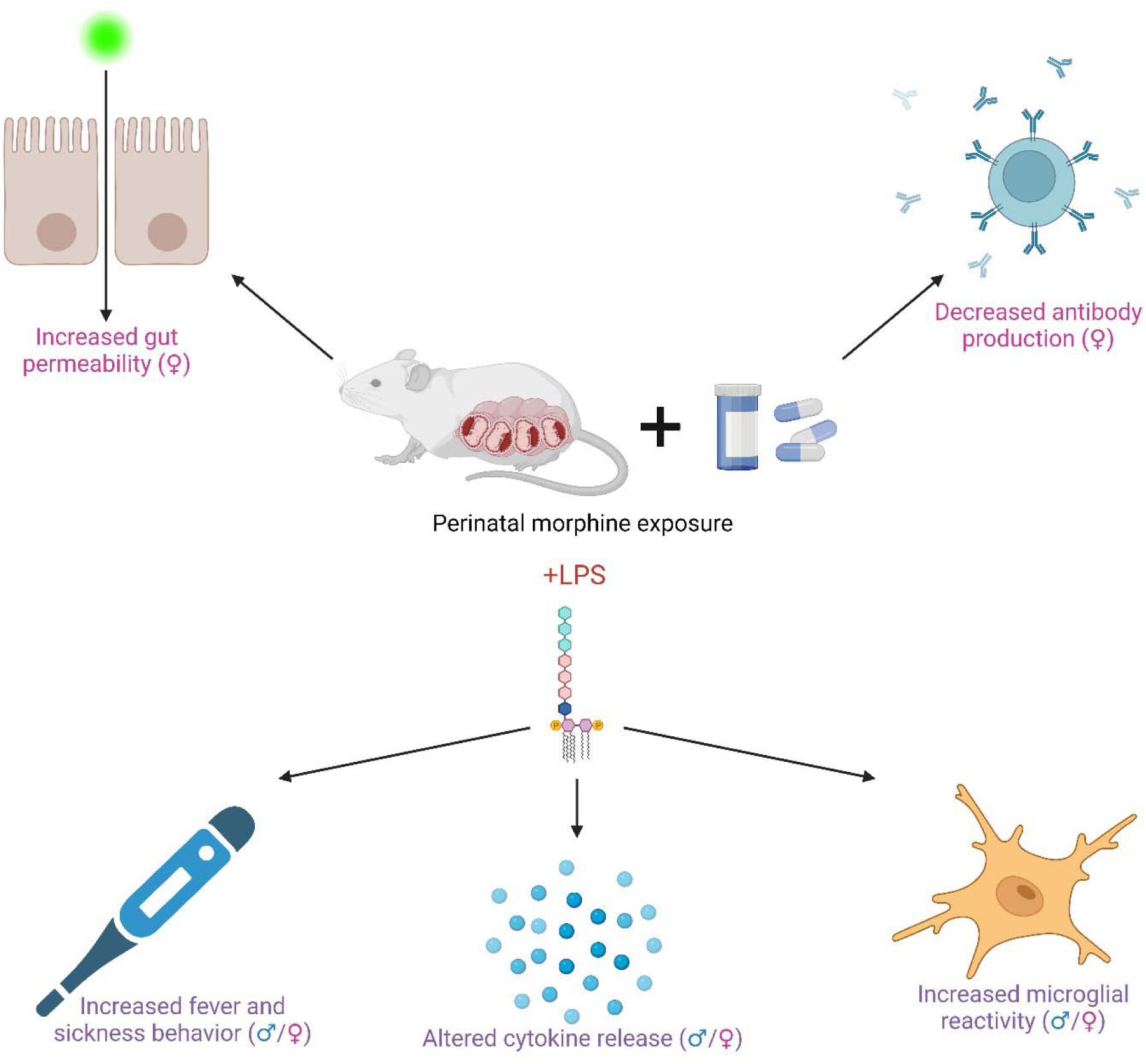
Summary of results. Created with BioRender.com.

## Acknowledgments

This work was supported by the National Institutes of Health (1RO1DA041529). The funding source was not involved in study design, data collection, analysis or interpretation, manuscript writing, or the decision to publish this article. Morphine sulfate was provided by the NIDA Drug Supply Program.

