## Supplemental for "Increased LPS-Induced Fever and Sickness Behavior in Adult Male and Female Rats Perinatally Exposed to Morphine"

Harder - Increased Fever in Rats Exposed to Morphine – Supplemental Table 1

|  | MOR M | | | MOR F | | |
| --- | --- | --- | --- | --- | --- | --- |
|  | Percent Change | Welch’s t value | p value | Percent Change | Welch’s t value | p value |
| Length | -18.0% | -3.16 | 0.003* | -22.5% | -3.67 | <0.001* |
| Area | -25.2% | -5.27 | <0.001* | -15.4 | -2.76 | 0.007* |
| Volume | -25.8% | -5.75 | <0.001* | -11.1% | -1.93 | 0.053 |
| Segments | -23.2% | -3.44 | <0.001* | -25.7% | -3.57 | <0.001* |
| Edges | -25.5% | -4.87 | <0.001* | -19.2% | -3.41 | 0.002* |
| Vertices | -25.5% | -4.87 | <0.001* | -19.1% | -3.41 | 0.002* |
| Sholl AUC | -12.3% | -2.10 | 0.034* | -25.2% | -4.23 | <0.001* |

**Supplemental Table 1.** Summary of alterations in microglial morphology in the entorhinal cortex. % represent percent change from sex-matched VEH controls. * = significant at p<0.05.
